# Cerebral blood flow measured with diffusing wave spectroscopy during anesthesia

**DOI:** 10.1101/438028

**Authors:** Markus Belau, Wolfgang Scheffer, Georg Maret

## Abstract

The adequate perfusion of the brain is of utmost importance where already short periods of hypoperfusion may lead to permanent damage. In order to increase patient safety the cerebral blood flow should be monitored in clinical settings during situations, such as anesthesia, where the perfusion might be disturbed. The cerebral blood flow is however not monitored on a routine basis during anesthesia. Diffusing wave spectroscopy is a relative novel optical method that non-invasively measures changes in cerebral blood flow. Here we report changes in cerebral blood flow associated with a delayed cardiac output, a change in isoflurane concentration and body temperature observed during general anesthesia with isoflurane in pigeons.

## 1 Introduction

The adequate perfusion of the brain is of utmost importance to avoid damage of the brain. In particular during anesthesia the cerebral autoregulation might be disturbed which may result in ischemia.^1^ However, during general anesthesia the cerebral blood flow (CBF) is not monitored on a routine basis.

The standard method to measure CBF in humans is transcranial Doppler (TCD) which however is limited to large arteries and the so called “ultrasonic window” where the ultrasound can reach large cerebral arteries.^2^ Other possibilities such as arterial spin labeling magnetic resonance imaging (MRI), xenon-enhanced computed tomography (XeCT), positron emission tomography or near infrared spectroscopy with indocyanine green injection either require large equipment and/or are invasive. In contrast, Diffusing wave spectroscopy (DWS)^3^ also known as diffuse correlation spectroscopy (DCS) measures the average local micro circulation and can be applied throughout the head.^4, 5^ Furthermore, it is relatively inexpensive, fast and portable and might greatly improve anesthetic management as well as chronic monitoring during or after events with disturbed perfusion such as stroke.^6–9^ It has been well validated against a variety of other modalities including arterial spin-labeled MRI,^10–12^ fluorescent microspheres,^13^ XeCT,^14^ Doppler Ultrasound,^15^ Laser Doppler^16–18^ and artificial perfusion of kidney^19^ (for an overview see e.g.^20^).

Here we demonstrate the potential of DWS in the monitoring of the cerebral blood flow during general anesthesia. In particular we demonstrate a linear relation of the microvascular cerebral blood flow with the relative delay of the cardiac output. Furthermore changes in cerebral blood flow associated with changes in the inspiratory isoflurane concentration and body temperature are shown.

One should note that the experiments where not specifically designed for the investigation of the above mentioned clinical effects but rather were incidental findings during another investigation.

Due to the significant changes they are nevertheless worthwhile to be presented and should be used as a starting point for a proper designed study on humans.

## 2 Methods

### 2.1 Diffusing Wave Spectroscopy

The principle of Diffusing wave spectroscopy for a semi-infinite geometry is schematically depicted in Fig. 1. Light with a long coherence length is used to illuminate the tissue and detected at a distance *ρ*. The light between source and detector is multiple scattered along different paths which are on average located in a banana shaped volume determining the spatial resolution and the penetration depth (about 1/2-1/3 of the source-detector separation). The light from the different paths interfere constructively and destructively at the surface and form a so called speckle pattern. If the scatterers move in time the paths and therefore the speckle pattern will change in time. The scatterer dynamics are mostly driven by the movement of red blood cells (except for large tissue motion during e.g. muscle contraction^21^). The light from one of these speckles is detected and the normalized intensity autocorrelation function *g*_2_(*τ*) is calculated. The autocorrelation function can then be fitted with the analytical solution of the diffusion equation to deduce the dynamics, namely the blood flow.

**Fig 1.**
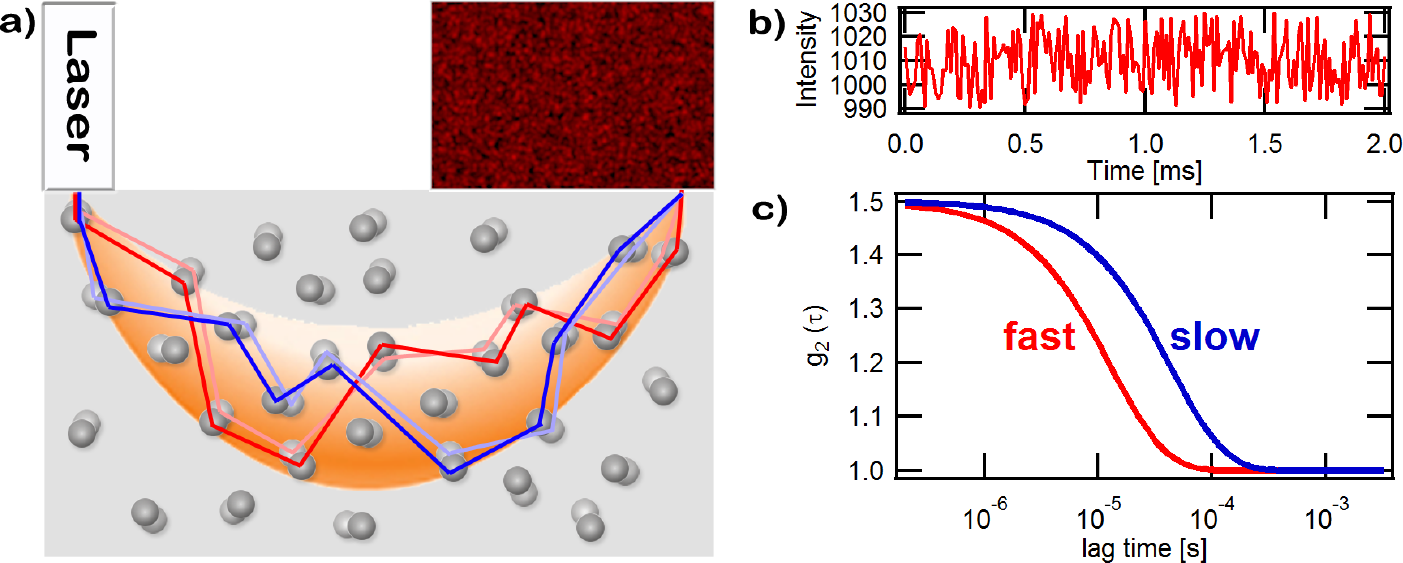
Principle of Diffusing Wave Spectroscopy. a) A laser with a long coherence length is used to illuminate the sample. The light is multiple scattered along different paths. The light from these different paths interfere constructively and destructively at the surface and build a so called speckle pattern. If the scatterers move in time the speckle pattern will also change with time. b) DWS measures the intensity fluctuations of one speckle. From the intensity fluctuations the autocorrelation function is calculated which measures how fast the intensity fluctuates in time. c) The normalized intensity autocorrelation function *g*_2_(*τ*) is shown. From the timescale of the decay and the shape one gets information about the dynamics (red: fast; blue: slow) and the type of motion of the scatterers (e.g. diffusive).

Measurements were performed with a fiber-multispeckle setup.^22^ Light from a diode laser (TOPTICA, TA100) operating at λ = 802nm is coupled to a multimode optical fiber to deliver light to the head. Multiple scattered light is detected by a fiber bundle consisting of 32 single-mode fibers (Schäfter+Kirchhoff SMC-780) which probes statistically equivalent but independent speckles. In order to perform the experiments in the presence of ambient light a bandpass filter (Semrock FF01-800/12-25) centered at 800nm is used in the receiver bundle. The light from each fiber is guided to an avalanche photodiode (APD; Perkin-Elmer SPCM-AQ4C). The output is connected to a custom build 32-channel multitau hardware correlator (correlator.com) which continuously computes normalized intensity autocorrelation function *g*_2_(*τ*) as a function of lag time *τ* with a integration time of 13ms. The bundle-average intensity autocorrelation function is calculated by an intensity weighted sum 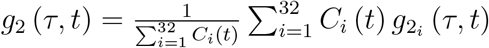, where *C*_*i*_(*t*) and 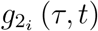 is the average intensity and intensity autocorrelation function of the i-th fiber, respectively. The normalized intensity autocorrelation function is fit by the solution of the diffusion model for a semi-infinite geometry:^23^

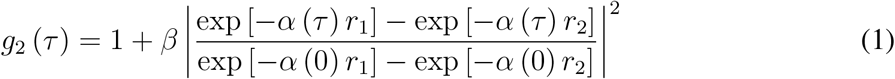

with the dynamic absorption coefficient 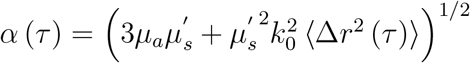 with 〈Δ*r*^2^(τ)〉 = 6*Dτ*, where *D* is the effective diffusion coefficient which is approximately proportional to blood flow.^5, 10–20^ We will therefore use the term cerebral blood flow (CBF) interchangeably for efeffective diffusion coefficient (D) throughout the paper. The quantities 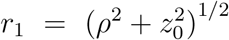 and *r*_2_ = [*ρ*^2^ + (*z*_0_ + 2*z*_*b*_]^2^)^1/2^ are related to the depth 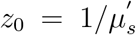 of the diffuse light source, the extrapolation length 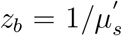 and the source detector separation *ρ*. The optical parameters have been fixed to the following values: absorption coefficient *μ*_*a*_ = 20m^−1^, reduced scattering coefficient 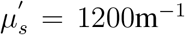, refractive index *n* = 1.4 and k_0_ = 2*πn/λ* denotes the wavevector. *β* is a coherence factor which accounts for the number of detected speckles and the detection optics and is about 0.5. The assumption of fixed optical parameters *μ*_*s*_ and *μ*_*a*_ may lead to some errors in the absolute diffusion coefficient between individuals, but since all our conclusions are only drawn from relative changes inter-individual changes only lead to negligible additions errors. Dynamic changes in the effective absorption coefficient due to changes in blood volume during e.g. a skipped heart beat are small (1-2% change in count rate) and since the effective absorption coefficient (20m^−1^) is much smaller than the effective scattering coefficient (1200m^−1^) the optical path length distribution is hardly affected at all. We therefore think that it is not necessary to measure the optical parameters for the proposed applications.

Measurements were performed at different positions of the head with source detector separations between 6.5mm and 20mm (average: 15.5 ± 2.9mm). We did not differentiate between intracerebral and extracerebral contribution of the blood flow by different source detector separations, but for the reported values we neither expect a localized change in only one of the compartments nor did we observe any effect in our data.

It has been shown in a number of studies that the fitted effective diffusion coefficient D is proportional to cerebral blood flow where a large value corresponds to a high blood flow and a low value to slow blood flow. The countrate is mainly given by changes in absorption and therefore is approximately inversely proportional to blood volume. Due to the high temporal resolution of our DWS system the heart beat and to an smaller extend the breathing are clearly visible (see Fig. 2). In order to analyze the data in more detail variations due to heart beat and breathing have been removed by an adaptive filtering approach where essentially a ‘typical’ heart beat and breathing temporal waveform have been detected and subtracted from the data to get rid of blood flow variations due to different times in the heart beat and breathing cycle.

The heart rate is determined from the diffusion coefficient by a custom written peak finding routine which identifies the peak position in the systole (gray lines in Fig. 2).

### 2.2 Narcosis

All reported experiments are part of a study where functional changes due to neural activity should be investigated. Therefore the experiments and narcosis protocol has not been optimized to study the effect of general anesthesia on the cerebral blood flow.

**Fig 2.**
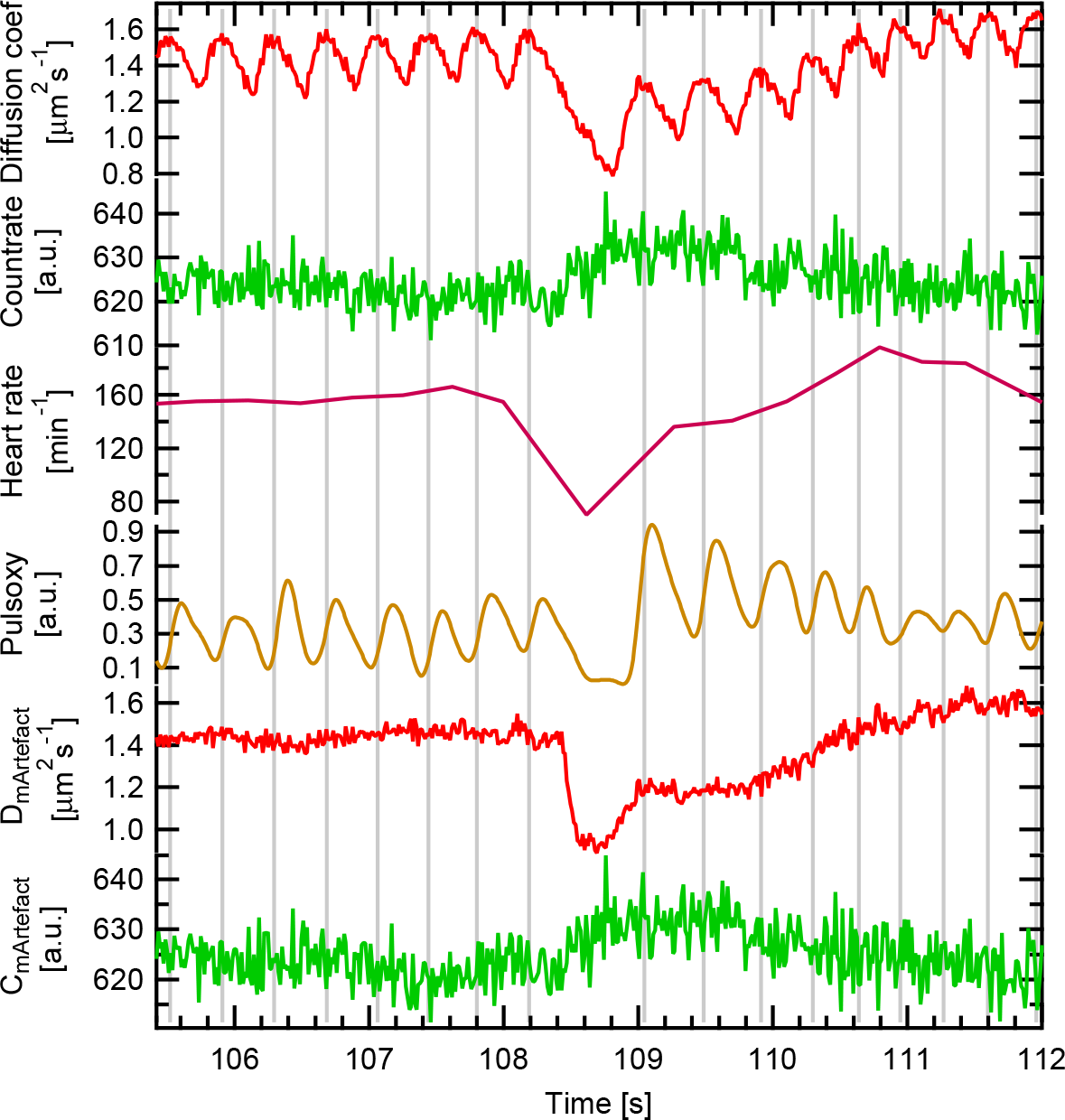
The figure shows a typical example for an arrhythmia where one heart beat has produced no cardiac output. The top graph represents the diffusion coefficient which is approximately proportional to cerebral blood flow, the count rate is approximately inversely proportional to blood volume. Heart beat has been determined from the peaks in the diffusion coefficient which are additionally indicated by gray vertical lines. The pulse oximeter signal is measured at the distal end of *M. gastrocnemius* and is shown in dark yellow. The lower two curves are the diffusion coefficient and the count rate after removal of variations due to heart beat and breathing.

The following narcosis protocol has been use: The pigeon is fasted for about 12h before anesthesia. Anesthesia is performed with the pigeon spontaneously breathing in an half open system where pure oxygen with a flow of 2l/min is used. The isoflurane concentration is adjusted by a isoflurane vaporizer (Eickemeyer, Isoflo) which can be adjusted in steps of 0.5%. The isoflurane concentration is adjusted by effect where anesthetic depth is accessed by the reflex score of Korbel et al.^24^ with an intended reflex score of 3 (surgical plane). Three anesthesia regimes have been used: 1) isoflurane as a single agent, 2) isoflurane with buprenorphine (0.5mg/kg i.m.), 3) isoflurane with midazolame (5mg/kg i.m.). For regime 1 and 2 the maintenance dose is about the same (≈3%) while in regime 3 the maintenance dose is about (≈2%). The numbers of the compositions of the different regimes will be given in the corresponding sections.

A pulse oximeter (SurgiVet V90041) continuously measures the saturation, heart rate and photo-plethysmogram at the distal *M. gastrocnemius*. Temperature is measured rectally with a digital thermometer (Voltcraft Multi-Thermometer DT-300) and tried to kept constant with a water perfused heating blanket. Nonetheless a drop in body temperature often occurred after induction of anesthesia (see section 3.3).

## 3 Results

### 3.1 Arrhythmia

Different types of arrhythmia have been observed during anesthesia. Here only the case with a delayed cardiac output will be described in detail. A typical time course of the delayed cardiac output is displayed in Fig. 2. The variations in blood flow during systole (high value of D - gray vertical lines) and diastole (low value of D) can be nicely seen. During a skipped heart beat the blood flow is decreased to about half it’s value compared to the regular blood flow. The pulse oximeter signal measured on the distal end of the left *M. gastrocnemius* shows the same effect albeit with a small delay. For the count rate the variations due to heart beat are significantly smaller, but can still be clearly identified in the Fourier transform (not shown). During the delayed cardiac output a small increase in count rate is observed. The curves D_mArtefact_ and C_mArtefact_ show the diffusion coefficient and count rate after adaptive filtering which removed variations due to heart beat and breathing.

To access the behavior more quantitatively in total *N* = 201 events with a delayed cardiac output (from 18 experimental days and 12 different pigeons, with 182, 1 and 18 artifacts in narcosis regime 1, 2 and 3 respectively) has been analyzed.

**Fig 3.**
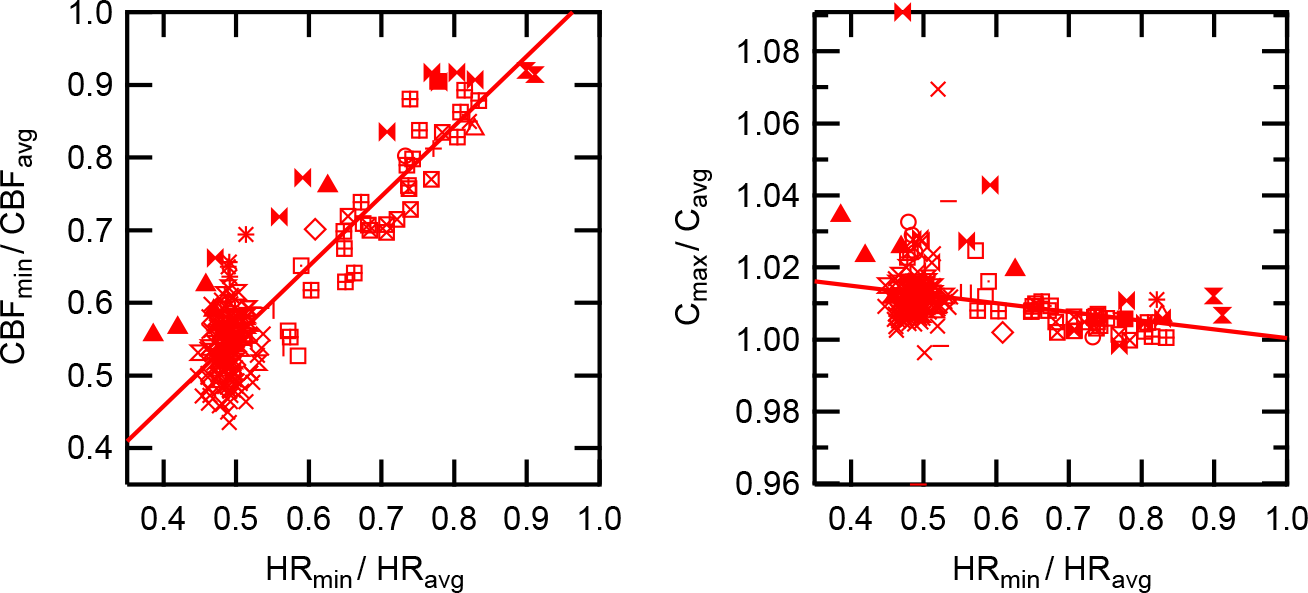
Left: relative cerebral blood flow as a function of delay of heart beat. Right: relative countrate vs relative heart rate. Different symbols correspond to results from different experimental days.

The dependence of the ratio of cerebral blood flow during delayed cardiac output and baseline cerebral blood flow is plotted as a function of the ratio from the heart rate during delay and baseline heart rate (Fig. 3). The blood flow shows a linear relation with the relative delay of the cardiac output (slope: 0:96 ± 0:04, r=0.90, *p* < 10^−10^). The count rate has a larger variability but also has a linear relation with the heart rate (slope: −0:024 ± 0:007, r=−0.248, *p* < 0:0005). If one normalizes the change during the delayed cardiac output to the average amplitude of the change during a normal heart beat one finds a slope of 3:41 ± 0:12 and −4:35 ± 2:53 for the blood flow and the count rate, respectively. The relative time of the diastole is 61±2% of the heart cycle. With a linear decay in flow this gives for a skipped heart beat a decay of 1:64 ± 0:05 times the heart beat amplitude which corresponds to a slope of 3:28 ± 0:11 which is consistent with the observed changes in blood flow and blood volume.

#### 3.1.1 Discussion

A linear correlation between heart rate and cerebral blood flow is found during the delayed cardiac output. The linear dependence of the cerebral blood flow can be simply understood (neglecting pulsatile effects for the moment): The cardiac output (CO) is given by stroke volume (SV) times the heart rate (HR)^25^

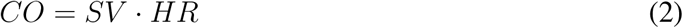

and the mean arterial blood pressure (MAP) is related to the systemic vascular resistance (SVR) and central venous pressure (CVP) by

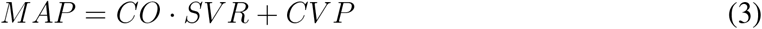

The cerebral blood flow (CBF) can then be calculated by Ohm’s law

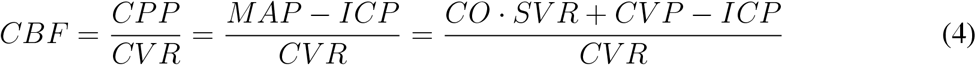

with CPP cerebral perfusion pressure and ICP intracranial pressure and CVR cerebral vascular resistance. Assuming that CVP and ICP are small and that the stroke volume and vascular resistance is approximately constant this reduces to

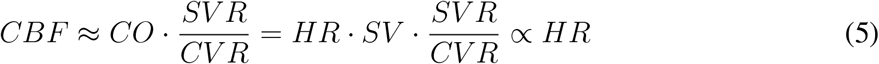

The slope of the decay during diastole is approximately proportional to the negative diffusion coefficient. This exponential behavior can be understood by the windkessel model of the blood vessels.^26^ Almost no effect due to wave reflection is seen in the data.

A delayed cardiac output both in the cerebral blood flow measured with DWS and in the Pulsoxy data measured on the distal *M. gastrocnemius* has been observed. The reason for the delayed cardiac output are most likely ventricular extrasystoles (premature ventricular contraction) with a compensatory pause, however no definite assignment can be made since no electrocardiogram has been recorded during the experiments. Arrhythmia, such as ventricular extrasystoles are quite common in anesthesia, but also occur in healthy people, and are considered to have a weak illness value and a benign prognosis. However the dramatic decrease in blood flow may have severe effects in situations such as during resuscitation and in critical ill patients where the maintenance of circulation is of utmost importance.

In addition blood flow monitoring is often performed with slow sampling rates (< 1Hz) which do not allow for the monitoring of variations during the heart cycle. In these measurements the effect of arrhythmia might be overlooked leading to wrong conclusions.

Note that the blood flow is not only affected during a delayed heart beat but also during an early heart beat (data not shown). The linear correlation of heart rate and blood flow allows to correct the data for these artifacts. This is of particular interest in situations, such as functional imaging, where one wants to distinguish between physiological changes and other effects.

### 3.2 Isoflurane concentration

During some measurements the isoflurane concentration has been changed by 0:5%. Figure 4 shows exemplary data of a measurement where the isoflurane concentration has been decreased from 2.5% at 50s to 2.0% and back to 2.5% at 250s. After the decrease of the isoflurane concentration an increase of about 8% is observed in the heart beat averaged blood flow which returned to the baseline value after return to baseline isoflurane concentration. Note that the change in blood flow during systole and diastole is quite a bit larger than the observed changes due to the isoflurane concentration (inset Fig. 4).

**Fig 4.**
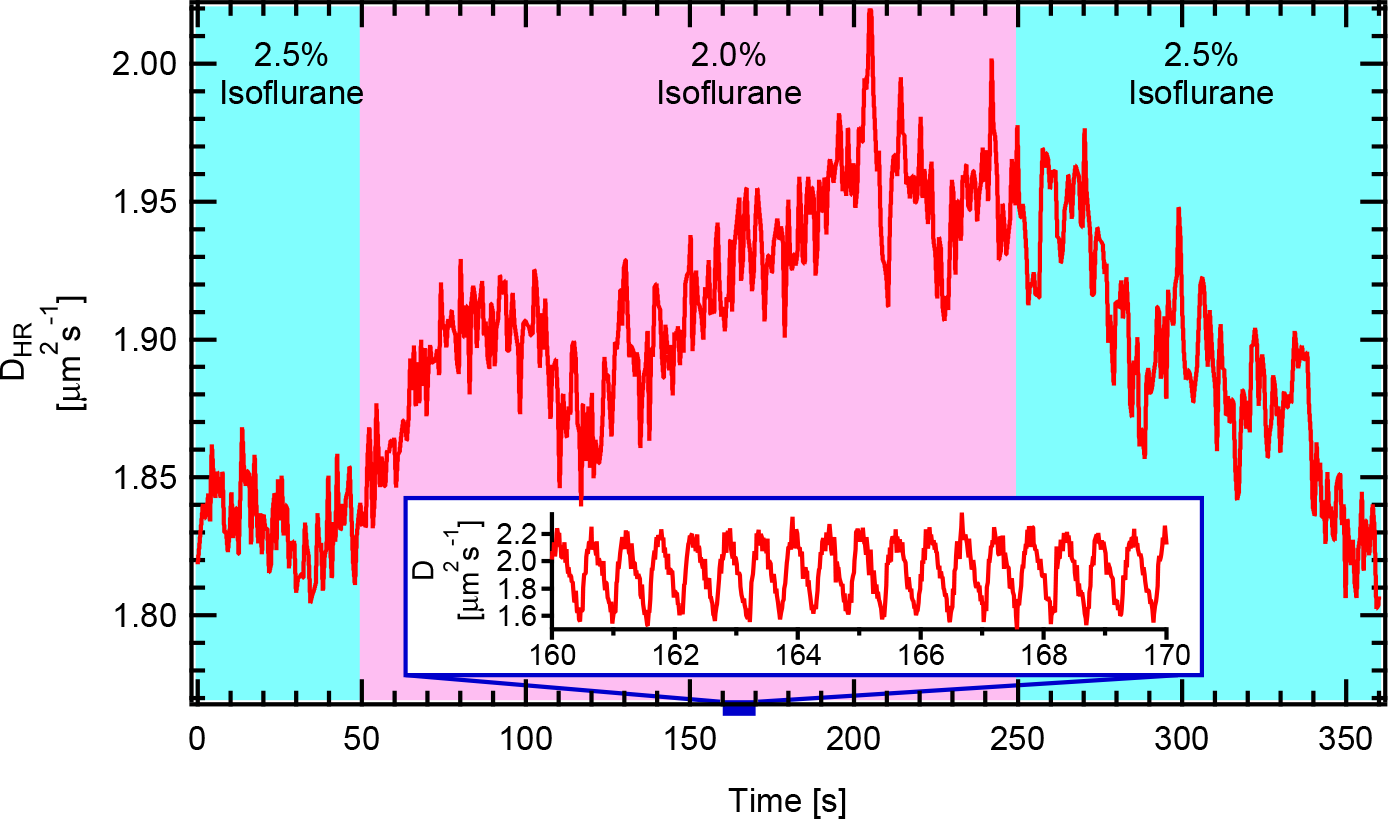
Change in the diffusion coefficient averaged over heart rate *D*_*H R*_ (cerebral blood flow) due to a variation in the isoflurane concentration. The inset shows the non-averaged diffusion coefficient.

**Fig 5.**
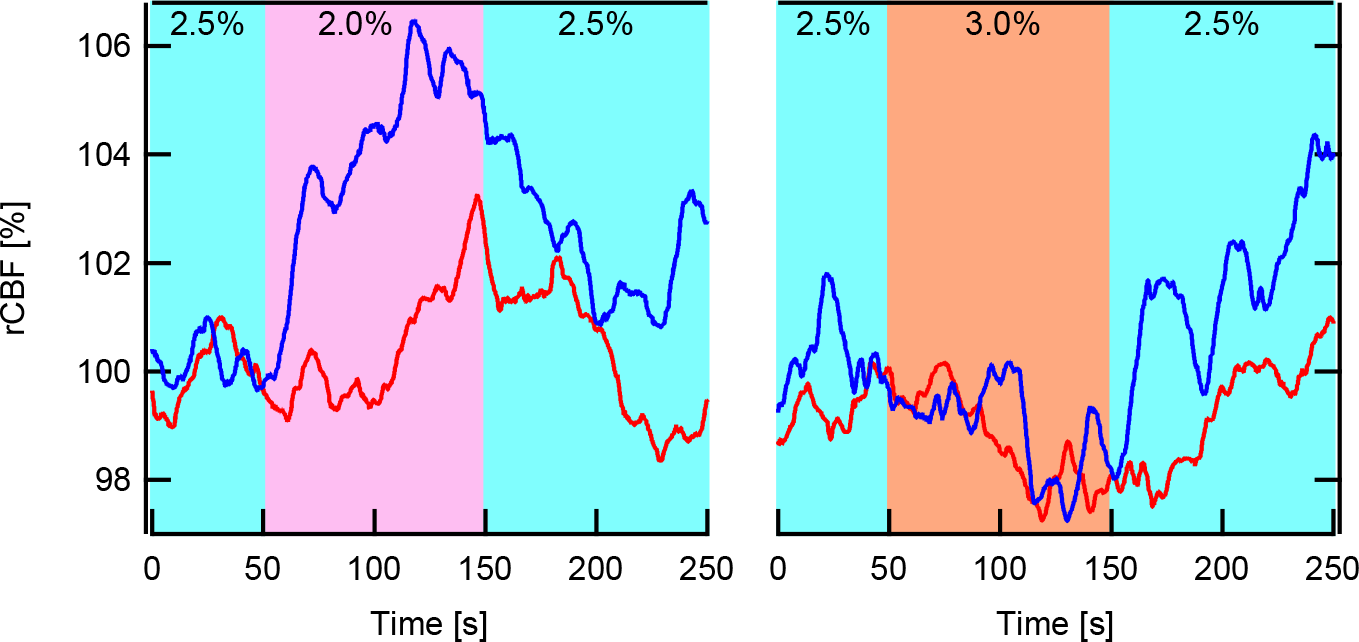
The figure exemplary displays the effect of a change in isoflurane concentration to the relative cerebral blood flow (rCBF) in two pigeons (red and blue). To make the data more clear the 10s median is shown. The maintenance dose was 2.5% (blue area) and has been decreased to 2.0% at 50s (pink area) and back to 2.5% at 150s followed by a measurement where the concentration has been increased to 3.0% at 50s (orange area) and back to 2.5% at 150s.

To understand the effect in more detail both a decrease and an increase of the isoflurane concentration has been performed for two pigeons (Fig. 5). Both pigeons have been in narcosis for about 40-45 minutes with a maintenance dose of 2.5%. For the decrease of isoflurane concentration one finds an increase and an opposite response for an increased concentration with a decrease in relative cerebral blood flow (rCBF). The amplitude and temporal course show some variations but with the same tendency.

In order to better understand the origin of the blood flow change the grand average of all measurements where the concentration has been changed during the measurement is calculated (Table 1 and Fig. 6). One finds that except for body temperature both groups have non significant different properties (however note the group size difference) but show an opposite behavior in the average cerebral blood flow.

**Table 1.** Group averages of physiological data from the measurements where isoflurane concentration has either been increased or decreased by 0.5%. Statistical analysis was performed with Student’s t-test and Wilcoxon-Mann-Whitney test. ns: not significant in both tests. *: *p* < 0:05 Student’s t-test. ^+^: *p* < 0:05 Wilcoxon-Mann-Whitney test.

|  | decrease | increase | significance |
| --- | --- | --- | --- |
| Number of Experiments | 15 | 4 |  |
| Buprenorphin | 2 | 0 |  |
| Isoflurane Concentration [%] | $2.63 \pm 0.3$ | $2.75 \pm 0.5$ | ns |
| Time in Narcosis [min] | $45 \pm 9$ | $39 \pm 8$ | ns |
| SpO2 [%] | $91 \pm 4$ | $92 \pm 4$ | ns |
| Heart rate [bpm] | $126 \pm 19$ | $131 \pm 18$ | ns |
| breathing rate [bpm] | $26 \pm 5$ | $23 \pm 4$ | ns |
| Weight [g] | $464 \pm 65$ | $495 \pm 80$ | ns |
| h Temperature [°C] | $40.8 \pm 1.0$ | $41.7 \pm 0.3$ | * |
| Source Detector Separation [mm] | $15 \pm 3$ | $17 \pm 2$ | ns |
| Average CBF change [%] | $1.6 \pm 5.3$ | $-2.2 \pm 1.6$ | *+ |

From Ohm’s law it follows that the change in blood flow might arise from a change in blood pressure or vascular resistance. Considering only the brain it follows that

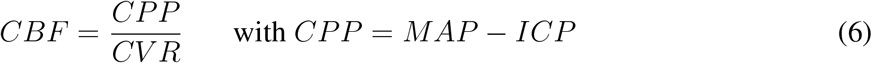

where CPP is the cerebral perfusion pressure, CVR cerebral vascular resistance, MAP the mean arterial blood pressure and ICP the intracranial pressure. Changes in CVR are often accessed with ultrasound by the Gosling Pulsatility index (PI):^27, 28^

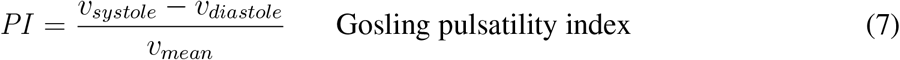

Here we use the effective diffusion coefficient to introduce a DWS pulsatility index in a similar manner:

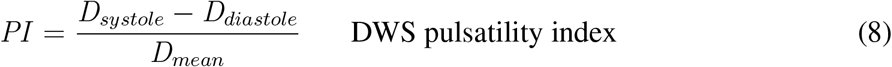

In literature often an linear dependence of PI with CVR is assumed, but there are situations where PI and CVR show inverse behavior^29^ (for a detailed discussion and a simple model for PI see^30, 31^). With the simple model one can nonetheless assume that |*PI*| ∝ |*CV R*| should be approximately valid. Therefore if there are no changes in the pulsatility index the cerebral vascular resistance should also be constant. Assuming furthermore that ICP is a lot smaller than MAP one gets

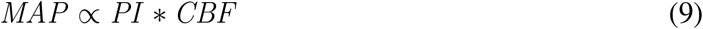

Figure 6 summarizes all measurements with a changed isoflurane concentration showing the percentage change of the normalized cerebral blood flow (ΔrCBF), the pulsatility index (ΔrPI) and the estimated mean arterial blood pressure (ΔrMAP). Solid lines show the average over all measurements, light dashed lines show individual measurements. It can be nicely seen that a decrease in isoflurane concentration leads on average to an increase in CBF and an increase of isoflurane to a decrease. It has to be noted that the data shows a great variability both in amplitude and time and even reverse effects can be seen in some instances. Nevertheless the difference is highly significant (*p* < 0:001) (Student’s t-test, Wilcoxon-Mann-Whitney test).

**Fig 6.**
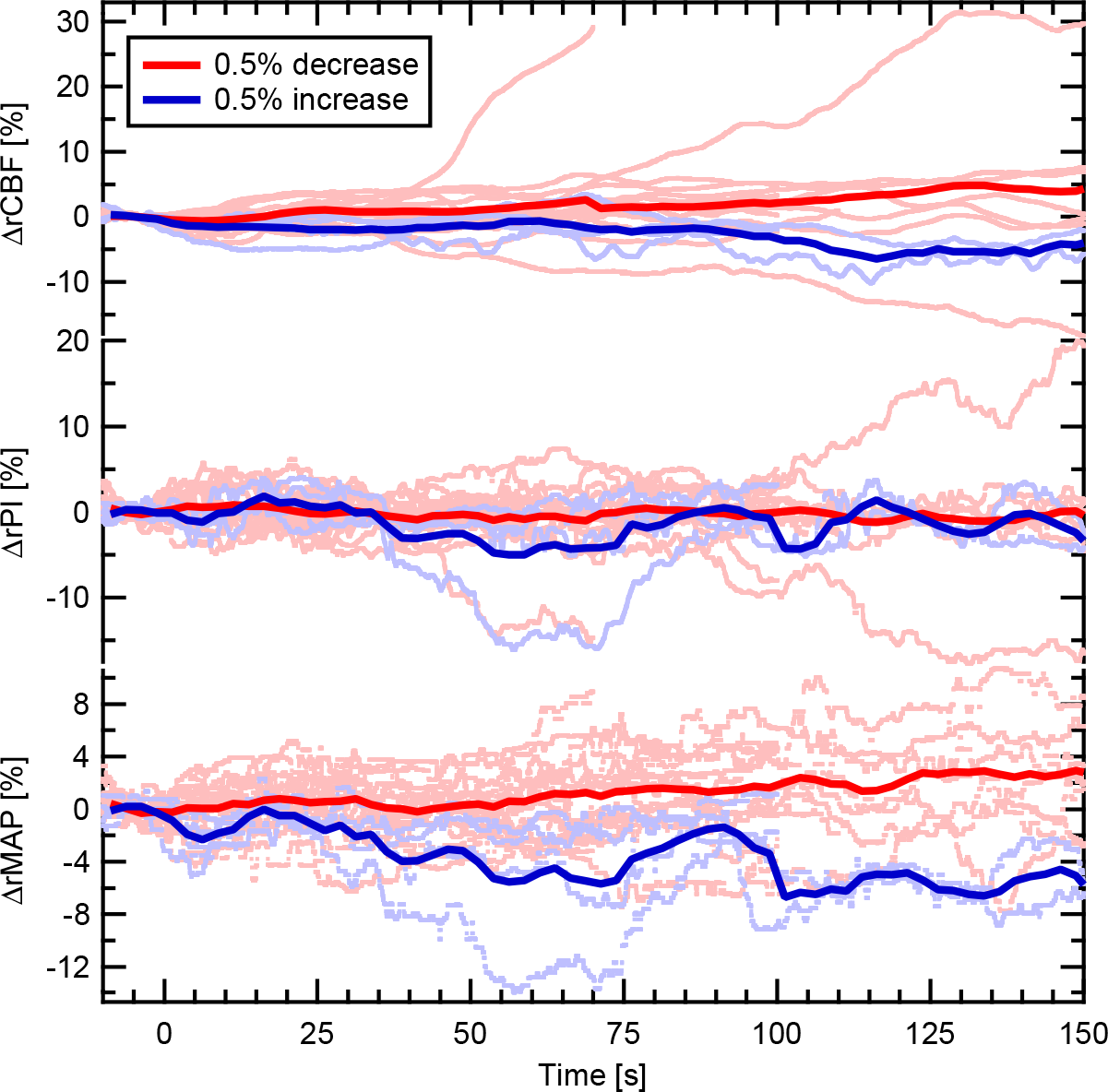
Average over all data where the isoflurane concentration has been decreased by 0.5% (red) and increased by 0.5% (blue) at 0s from the maintenance dose (average 2.66%. (2.0%(1), 2.5%(12), 3.0%(5), 3.5%(1))). The solid lines indicate the average over all measurements and the light dotted lines show all data which were used for the averages. All data has been normalized to the average value in the range −10s to 0s. ΔrCBF denotes the percentage change in cerebral blood flow, ΔrPI the change in the pulsatility index, ΔrMAP the estimated change in mean arterial blood pressure.

In contrast the pulsatility index is nearly constant over time. Some stronger variation can be seen in the case of increased concentration which are due to additional artifacts in the data (e.g. ventricular extrasystole with compensatory pause which could not be fully compensated in data analysis). This results in some errors in the determination of the pulsatility index and due to the smaller number of experimental data (n=15 decrease; n=4 increase) for the increase this does not fully average out. The change in the estimated mean arterial blood pressure shows a similar behavior to cerebral blood flow, although the increase in MAP during decreased isoflurane concentration is smaller and the decrease during increased isoflurane concentration is larger compared to the change in CBF. The difference between increased and decreased data is however approximately constant in both MAP and CBF.

#### 3.2.1 Discussion

The observed response to a change in isoflurane concentration is somehow surprising since it is reported in literature that an increase in isoflurane concentration is associated with a cerebral vasodilation which results in an increase of blood flow.^32–34^ However this response is reported for concentrations usually used in humans (1-2%) which are considerably lower than the concentrations used in pigeons and applied in this study (2-3%). A likely explanation is that at the high concentrations used in pigeons the cerebral autoregulation might already be disturbed. In this regime the dominant effect is a change in mean arterial blood pressure which shows an increase during decreased isoflurane concentration and vice versa. Additional proof for this hypothesis is given by the fact that the pulsatility is nearly unaffected by changes of isoflurane concentration indicating that cerebrovascular resistance is constant in the range of the applied changes. However one has to note the limitations of these finding: First, many important physiological parameters such as blood pressure, end expiratory CO_2_ concentration, end tidal isoflurane concentration, narcotic depth (from electroencephalography or bispectral index), metabolic rate, ecg etc. have not been measured. In particular CO_2_ and in a smaller amount O_2_ also influence the vascular diameter and thereby the blood flow. In order to fully assign the effect to the concentration of the inhalation anesthetic, the other gas concentrations should be kept constant. Monitoring of the end tidal isoflurane concentration would allow to monitor the wash-in and wash-out times and identify the time when equilibrium is reached. Monitoring of blood pressure could confirm the assumed relationship and would allow for a more thorough analysis. In addition monitoring the pulsatility index both with DWS and TCD would allow to provide a quantitative relationship between both modalities. Second, one should use a standardized study protocol with equal group sizes to infer more meaningful quantitative values.

Nevertheless this preliminary work already demonstrates the great potential of DWS in anesthetic monitoring and anesthesia research. In particular it shows that, even though average values might be similar, the monitoring of each individual is very important because of the great variability in the response to a change in isoflurane concentration.

### 3.3 Body temperature

The body temperature is an important factor in narcosis because a loss of temperature is associated with a higher number of critical incidents but may also be used as a therapeutic tool during certain interventions to decrease the metabolic rate and thereby is protective to the tissue. Here we report the effect of body temperature on cerebral blood flow during isoflurane anesthesia in pigeons.

**Fig 7.**
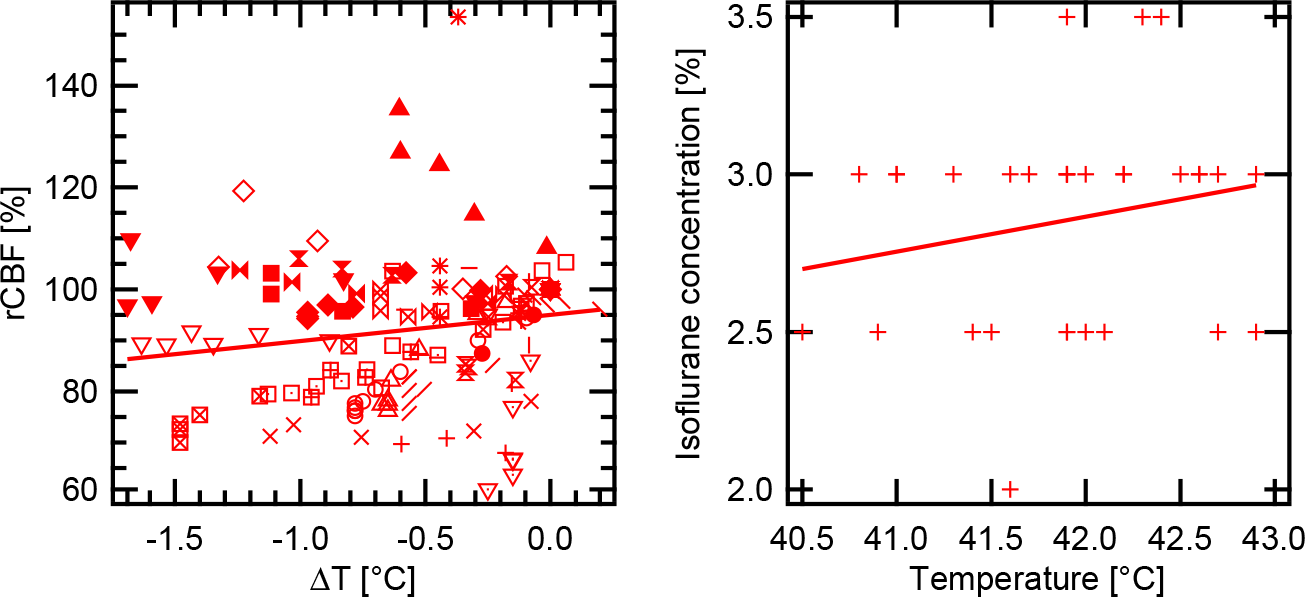
Left figure shows relative diffusion coefficient normalized to the time when the first temperature was measured versus the temperature difference relative to the first measured temperature. Different symbols correspond to different experiments. The right figure shows the maintenance dose of isoflurane versus the first measured temperature. The straight line represents a linear regression of the data with a slope of 0:05 ± 0:02 and 0:11 ± 0:10% per °C for the left and right figure, respectively.

Figure 7 left shows the data from all measurements with isoflurane as a single agent against temperature difference relative to the first measured temperature. One finds on average that the rCBF is significantly reduced by about 5% per °C of decreased temperature (n=168, *r* = 0:178, *p* < 0:021).

The maintenance dose of isoflurane is shown as a function of the first measured temperature (Fig. 7 right). Although the correlation is not significant (n=31, *r* = 0:203, p=0.27) the slope in the linear regression of the data demonstrates that the first measured temperature maybe useful as a predictor for the maintenance dose.

No correlation can be found with weight (n=31, r=−0.085, not significant) and age (n=31,r=−0.089, not significant).

#### 3.3.1 Discussion

The measured data demonstrates a significant correlation of the cerebral blood flow during isoflurane anesthesia with body temperature. One has to note that not only the body temperature has been changed during the experiments, but also functional stimulation with light and changes in the isoflurane concentration have been applied in some of the measurements. No corrections for the correspondent effects have been performed. In addition one finds a tendency, albeit not significant, to require a higher maintenance dose if the first measured temperature is higher. One has to note that the first temperature has not been measured before induction of anesthesia but shortly after loss of consciousness (9.5 ± 3.3 min, range 5-20 min) where already a small drop in body temperature might have occurred and that the focus of the experiments where not on setting up the minimal required maintenance dose, which can also only be set up coarsely in steps of 0.5%. With an optimized study protocol with an adequate group size the results are likely to be significant.

One explanation might be that the increased body temperature leads to higher metabolism and therefore requires a higher isoflurane concentration. The increased metabolism will certainly have an effect, but the data can nearly completely be explained with the temperature solubility of isoflurane. Lockwood et al.^35^ reported that the solubility increases by 5.4% (compared to the value at 37°C) for each degree that equilibration temperature was reduced. Note that no data is available for temperatures above 40°C. Also for a larger temperature range the relation is rather logarithmic than linear.^36–38^

Nevertheless reported changes are in good agreement with the observed data with a slope of 5 ± 2% per °C of the relative blood flow and a change relative to 2.5% isoflurane of 4 ± 4% per °C. This is also in good agreement with data of Liu et al.^39^ who investigated the effect of hypothermia on the MAC in children and found a reduction of 5.1% per °C reduced body temperature.

## 4 Conclusion

Diffusing wave spectroscopy is a non-invasive, relative inexpensive, fast, portable method to continuously measure cerebral blood flow during general anesthesia. It is demonstrated that the blood flow shows a linear relation with the delay of the heart beat and thus is reduced by 50% during a skipped heart beat. Most of the modalities measuring cerebral blood flow have sampling rates which do not allow for the measurement of blood flow during individual heart beats. Since ventricular extrasystoles are quite common, particularly during anesthesia, this may lead to wrong conclusions drawn from the data of these modalities if the data is not corrected for these effects or data during arrhythmia is not excluded from analysis. The strong dependence of the cerebral blood flow on heart rate highlights the necessity to also measure the heart rate during functional studies where hemodynamic responses are typically small and one wants to discriminate between functional and physiological changes.

Even with the limitation of the presented preliminary results it is clearly shown that changes associated with inspired isoflurane concentration and body temperature can be measured. The deviation of single measurements from the group average behavior demonstrates the need to monitor cerebral blood flow for each individual to increase patient safety. We therefore suggest to validate the results in humans with a well designed study protocol and introduce DWS as standard monitoring during general anesthesia. This would also allow to identify correlates of blood flow changes associated with disease conditions.

## Acknowledgments

The research was supported by the Deutsche Forschungsgemeinschaft (DFG) with the Reinhart Koselleck project MA817/9-1 “Imaging of the brain response to magnetoreception”.

All experiments were performed in agreement with the regulations of the German Animal Welfare Act (TierSchG) and have been approved by the regional committee of animal welfare (DWS 35-9185.81 G-09 43, DWS 35-9185.81G-13 111).

